# Effect of terminal phosphate groups on collisional dissociation of RNA oligonucleotide anions

**DOI:** 10.1101/2024.03.05.583607

**Authors:** Mei-Qing Zuo, Ge Song, Meng-Qiu Dong, Rui-Xiang Sun

**Author notes:** **Correspondence** Meng‐Qiu Dong, Rui-Xiang Sun.

## Abstract

An increasing need of mass spectrometric analysis of RNA molecules calls for a better understanding of their gas-phase fragmentation behaviors. In this study, we investigated the effect of terminal phosphate groups on the fragmentation spectra of RNA oligonucleotides (oligos) using high-resolution mass spectrometry (MS). Negative-ion mode collision-induced dissociation (CID) and higher-energy collisional dissociation (HCD) were carried out on RNA oligos containing a terminal phosphate group on either or both ends, or neither. We find that terminal phosphate groups affect the fragmentation behavior of RNA oligos in a way that depends on the precursor charge state and the oligo length. Specifically, for precursor ions of RNA oligos of the same sequence, those with 5’- or 3’-phosphate, or both, have a higher charge state distribution and lose upon CID or HCD the phosphate group(s) in the form of a neutral (H_3_PO_4_ or HPO_3_) or an anion ([H_2_PO_4_]^-^ or [PO_3_]^-^). Such neutral or charged loss is most conspicuous for precursor ions of an intermediate charge state, e.g. 3^-^ for 4-nt oligos or 4^-^ and 5^-^ for 8-nt oligos. This decreases the intensity of sequencing ions (*a-, a-B, b-, c-, d-, w-, x-, y-, z-*ions), hence unfavorable for sequencing by CID or HCD. Removal of terminal phosphate groups by calf intestinal alkaline phosphatase improved MS analysis of RNA oligos. Additionally, the intensity of a fragment ion at *m/z* 158.925, which we have identified as a dehydrated pyrophosphate anion ([HP_2_O_6_]^-^), is markedly increased by the presence of a terminal phosphate group. These findings expand the knowledge base necessary for software development for MS analysis of RNA.

## 1. Introduction

Ribonucleic acid (RNA) is a linear polymer of adenine (A), uracil (U), cytosine (C), and guanine (G) ribonucleotides. RNA molecules carry out diverse biological functions in the cell. For example, messenger RNAs (mRNAs) carry genetic information from DNA to ribosomes. Ribosomal RNAs (rRNAs), along with ribosomal proteins and transfer RNAs (tRNAs), are responsible for synthesizing proteins according to the genetic information encoded by mRNAs. Small nuclear RNAs (snRNAs) are required for splicing of pre-mRNAs. As a representative of short, regulating RNAs, microRNAs (miRNAs) can control the transcription or translation of many target genes or mRNAs.^1,2^

RNAs also serve as research reagents or therapeutics. Antisense oligonucleotides, miRNAs, and small interfering RNAs (siRNAs),are examples in this category. In the recent Covid-19 pandemic, mRNA vaccines provided vital protection to more than five billion people.^3^

Functional diversity of RNA molecules is accompanied by diversity in length, sequence, and modification. The length of mRNAs varies from hundreds to tens of thousands of nucleotides (nt), whereas that of miRNAs lies in a narrow range of 19∼25 nt, or 22 nt on average. RNA sequence diversity is expanded by modifications, which have amounted to >170 types, including 2’-O-methylation, pseudouridylation, and m^6^A methylation, among others.^4^ Sequence and modification together determine the function and stability of an RNA molecule. As RNA research flourishes, the demand for RNA analytical methods increases.

Mass spectrometric analysis of RNAs can be traced back to 1960s for analysis of monomeric nucleic acids.^5,6^ Over the years, different ionization methods and fragmentation methods were explored,^7-13^ and application of MS in RNA research included identification of RNA sequences,^14,15^ modifications,^16,17^ and single-nucleotide polymorphisms,^18^ as well as quality examination of synthetic RNAs.^19^

Among the different fragmentation methods, CID and HCD are used more often than others for MS analysis of RNA oligonucleotides (oligos). In CID/HCD spectra, the predominant fragment ions of RNA oligos are *c*- and *y*-ions, arising from cleavage of the 5’ P-O bond.^20^ Of all sequencing ions (*a, a-B, b, c, d, w, x, y, z*) resulted from the breaking of the phosphodiester bonds, the five most abundant ones are *y, c, w, a-B*, and *a* ions.^21^ Factors that affect the fragmentation behaviors of RNA oligos include the precursor charge state, fragmentation energy, metal ion adduction, nucleotide modification, and the presence or absence of 3’-terminal phosphate.^15,17,20^

In this study, we investigated how the 5’- and the 3’-phosphate groups might affect the fragmentation behaviors of RNA oligos in the negative-ion mode. We compared the CID and HCD spectra of two sets of RNA oligos. Each set consisted of four oligos of the same sequence, but of different terminal groups, that is, with a hydroxyl group at both 5’- and 3’-end, or with a phosphate group attached to either or both ends. We found that the terminal phosphate group(s) up-shifted the precursor charge state and affected the CID and HCD spectra of RNA oligos similarly and profoundly. Specifically, they caused extensive loss of phosphoric acid neutral and metaphosphoric acid anion from precursors and diminished the intensity of sequencing ions. The presence of a terminal phosphate group enhanced markedly the intensity of *m/z* 158.925, which we identified as a dehydrated pyrophosphate anion, [HP_2_O_6_]^-^. Effective removal of the terminal phosphate groups by calf intestinal alkaline phosphatase was demonstrated.

## 2. Materials and methods

### 2.1. Materials and sample preparation

Eight RNA oligos with two sequences (GUCA and AUCGAUCG) and four types of terminal-group combinations (from 5’ to 3’: HO-X-OH, HO-X-P, P-X-OH, P-X-P, ‘X’ denotes one of the two sequences and ‘P’ is the phosphate group.) were synthesized by Biolino Acid Technology (Tianjin, China). The detailed information of these RNA oligos is provided in Supplemental Table S1. For direct infusion MS analysis, RNA oligos were dissolved in water to make 0.15 mM solutions (measured by NanoDrop 1000 Spectrophotometer, Thermo Fisher Scientific) and then diluted to 5 μM in 7.5 mM hexylamine (Energy chemical, Shanghai, China), 50% acetonitrile (Fisher Scientific) before being sprayed into mass spectrometry using a syringe pump. For LC-MS/MS analysis, 250 pmol of each RNA oligo was treated with 10 units calf intestinal alkaline phosphatase (CIP, New England Biolabs Inc., M0290S) in 37°C for 30 minutes. Then, the digests were desalted using ZipTip Piette Tips (Merck Millipore) following the manufacturer’s protocol before LC-MS/MS analysis. 250 pmol of each RNA oligo without CIP and desalting treatment was also conducted LC-MS/MS analysis as the negative control.

### 2.2. Mass spectrometric data acquisition

All experiments were performed on an Orbitrap Fusion™ Lumos™ Tribrid™ Mass Spectrometer (Thermo Fisher Scientific) in the negative-ion mode. For direct infusion analysis, the sample solutions were sprayed into the electrospray source at a flow rate of 2 μl/min by a syringe pump. The ion spray voltage was -2.5kV. The MS1 mass range was 150∼1500 *m/z* for GUCA and 180∼2700 *m/z* for AUCGAUCG. An inclusion list including all theoretical precursor *m/z* values from charge 1^-^ to charge 9^-^ for each RNA oligo was used. The fragmentation method was CID or HCD. The normalized collisional energy (NCE) was varied from 10% to 50%, with a step of 5%. For MS/MS, auto mass range modes were selected for all samples. The AGC target was 1E6 for MS1 and 1E5 for MS2. The maximum injection time was automatically adjusted by the system for both MS1 and MS2. Orbitrap was used as the mass analyzer for both MS1 and MS2 at a resolution of 120k. The isolation window was 0.4 *m/z* to reduce the proportion of mixed spectra. LC-MS/MS analysis was performed on an Easy-nLC 1200 UHPLC (Thermo Fisher Scientific) coupled to the Orbitrap Fusion™ Lumos™ Tribrid™ Mass Spectrometer. RNA samples were loaded and separated on a precolumn (75 μm inner diameter, 7 cm long, packed with Xbridge BEH C18 2.5 μm beads from Waters Corporation) and sprayed by an empty column (75 μm inner diameter, 3 cm long) with an in-housed pulled 5 μm tip. The column end fittings (Valco Instruments Company Incorporated, C-NEF.5XFPK.35S1) were used for the connection of liquid chromatography, the precolumn and the empty column to avoid the dislodge or fracture of the in situ cast silicate-based frits.^22^ A linear gradient made with buffer A (7.5 mM triethylamine, Thermo Fisher Scientific, 25108) and buffer B (80% acetonitrile and 7.5 mM triethylamine) as follows: 0-22 min, 0-30% B; 22-23 min, 30-90% B; 23-27 min, 90% B; 27-28 min, 95-0% B; 28-30 min, 0% B; 30-45 min, 0% B.

The flow rate was 350 nl/min. Cycle time was set to 3s. The resolution was 120k for MS1 and 60k for MS2.

### 2.3. Data analysis

Data analysis was conducted using in-house python and MATLAB scripts. Theoretical *m/z* values of *a-, a-B,b-, c-, d-, w-, x-, y-, z-* ions, internal fragments, and the precursors with a single nucleobase loss were calculated and matched with the observed peaks. For RNA oligos with a terminal phosphate group(s), the precursors with the loss of phosphoric acid or metaphosphoric acid anion were also calculated and matched with the observed peaks. Cyanate anion (NCO^-^) loss was not included here since the terminal pyrimidine bases were not contained in all oligos we used^21,23^. The observed monoisotopic peak was assigned to a theoretical ion if its mass tolerance is within 10 ppm. For CID spectra, NCE 35% was chosen as the representative energy because our data analysis (not shown) indicated that the sequencing ions are optimal at NCE 35% in terms of their total intensity. For HCD spectra, NCE 40% was selected to study the pattern of low-mass ions (within *m/z* 100 to 220).

## 3. Results and discussion

### 3.1. A terminal phosphate group tends to impart higher negative charges to precursor ions of RNA oligos

We analyzed two sets of RNA oligos (Supplemental Table S1), one with the sequence GUCA and the other AUCGAUCG. Within each set, the four oligos differ only at the 5’ or 3’ end, which is either –OH or –H_2_PO_4_. RNA oligos at a concentration of 5 μM each were introduced into a Q-Orbitrap instrument by electrospray ionization and analyzed in the negative-ion mode. As shown in Figure 1, the presence of 5’- or 3’-phosphate, or both, up-shifted the charge state distribution of the RNA precursor ions. The most abundant charge state of HO-GUCA-OH is 2^-^ (57.0%), followed by 3^-^ (36.0%), 1^-^ (6.2%), and 4^-^ (0.8%). The addition of one or two terminal phosphate groups leads to an abundance decrease of the 1^-^ and 2^-^ precursors, an abundance increase of the 4^-^ precursor, and even the appearance of a 5^-^ precursor in the case of P-GUCA-P where P denotes phosphate. The original MS1 signal intensities were shown in Supplemental Figure S1. Similarly, for AUCGAUCG, adding one or two terminal phosphate groups shifts the highest charge state of precursor ions from 6^-^ to 7^-^, and the charge state of highest abundance from 2^-^ to 5^-^ (Figure 1). Taken together, a terminal phosphate group imparts higher negative charges to precursor ions of RNA oligos.

**Figure 1.**
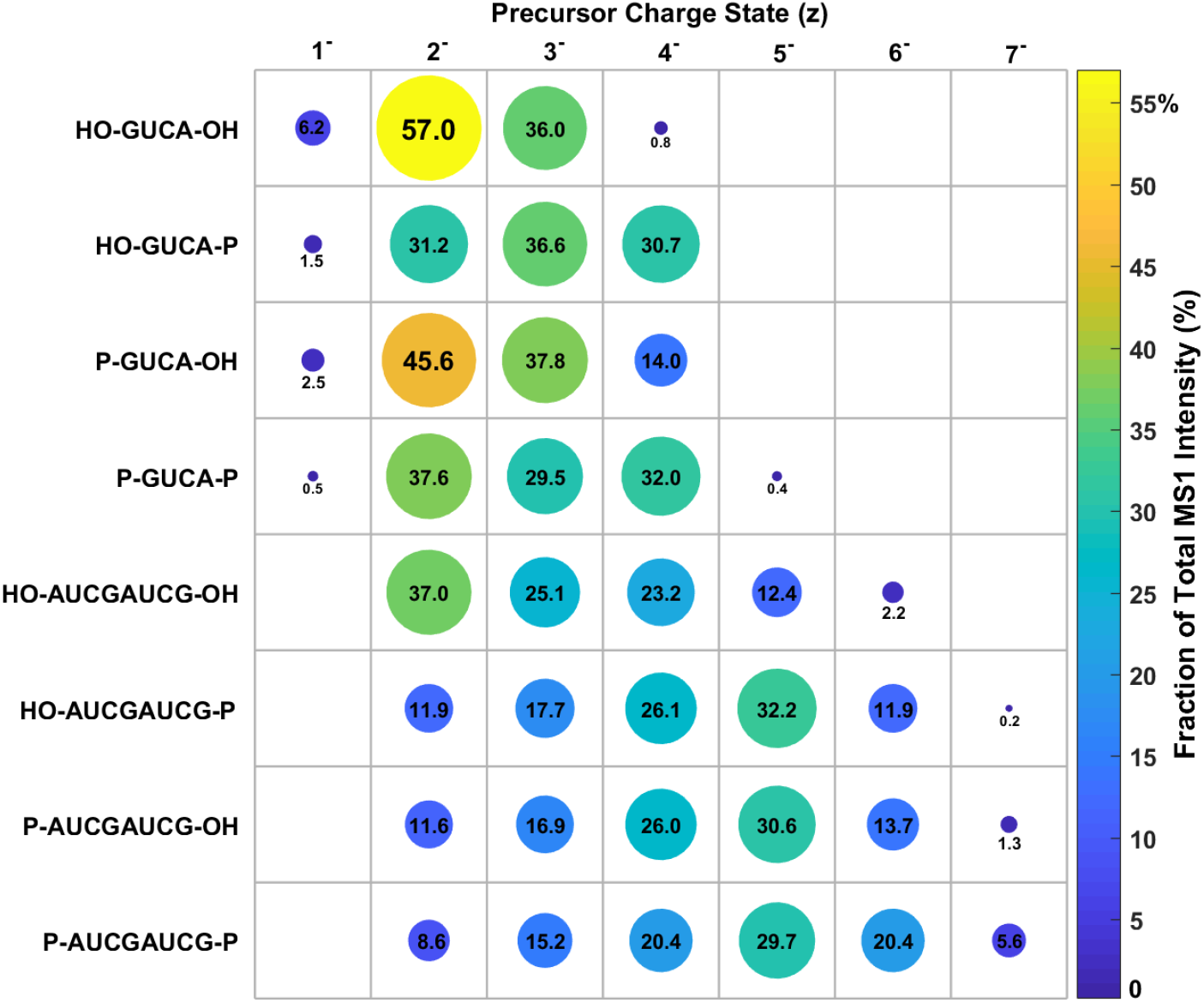
The charge state distributions of eight RNA oligonucleotides. The number inside or below each circle is the percentage of the MS1 intensity of the indicated precursor ion divided by all the precursor ions observed for each RNA oligo. A higher percentage is denoted by a larger diameter and a warmer color.

### 3.2. A terminal phosphate group(s) bestows a predominant loss of phosphoric acid or metaphosphoric acid from RNA oligo precursors with medium charges

A terminal phosphate alters not only the precursor charge state distribution, but also the fragmentation spectra of RNA oligos. Figure 2 shows the CID spectra from four GUCA oligos of charge 2^-^, fragmented under NCE 35%. The peaks from precursor ions with a neutral or charged loss of phosphoric acid ([M-H_3_PO_4_] or [M-H_2_PO_4_^-^]) or metaphosphoric acid ([M-HPO_3_] or [M-PO_3_^-^]) are highlighted in red and annotated in Figures 2b, 2c, and 2d. Such peaks were not observed in CID spectra of HO-GUCA-OH (Figure 2a). For the other three GUCA oligos with one or two terminal phosphate groups, the [M-H_3_PO_4_] peaks were the base peaks in their CID spectra (Figures 2b, 2c, and 2d). A consequence of such predominant (meta)phosphoric acid loss (PL) is the lower abundance of sequencing ions (compare Figures 2b∼2d with 2a). Similar results were obtained from the AUCGAUCG oligos of charge 4^-^ (Figure 3). The difference is that the base peaks from three AUCGAUCG oligos with one or two terminal phosphate group(s) are all [M-PO_3_^-^].

**Figure 2.**
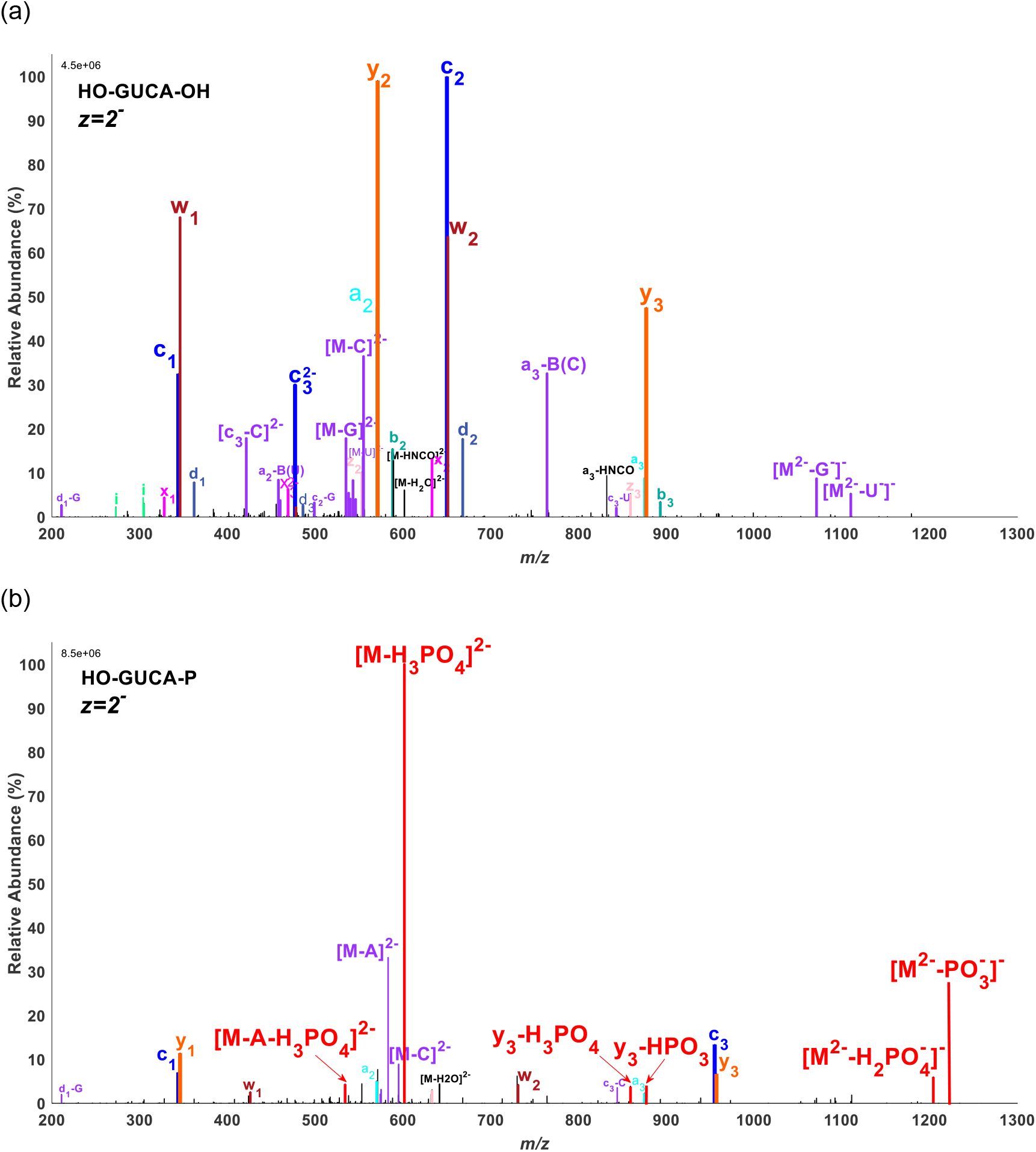
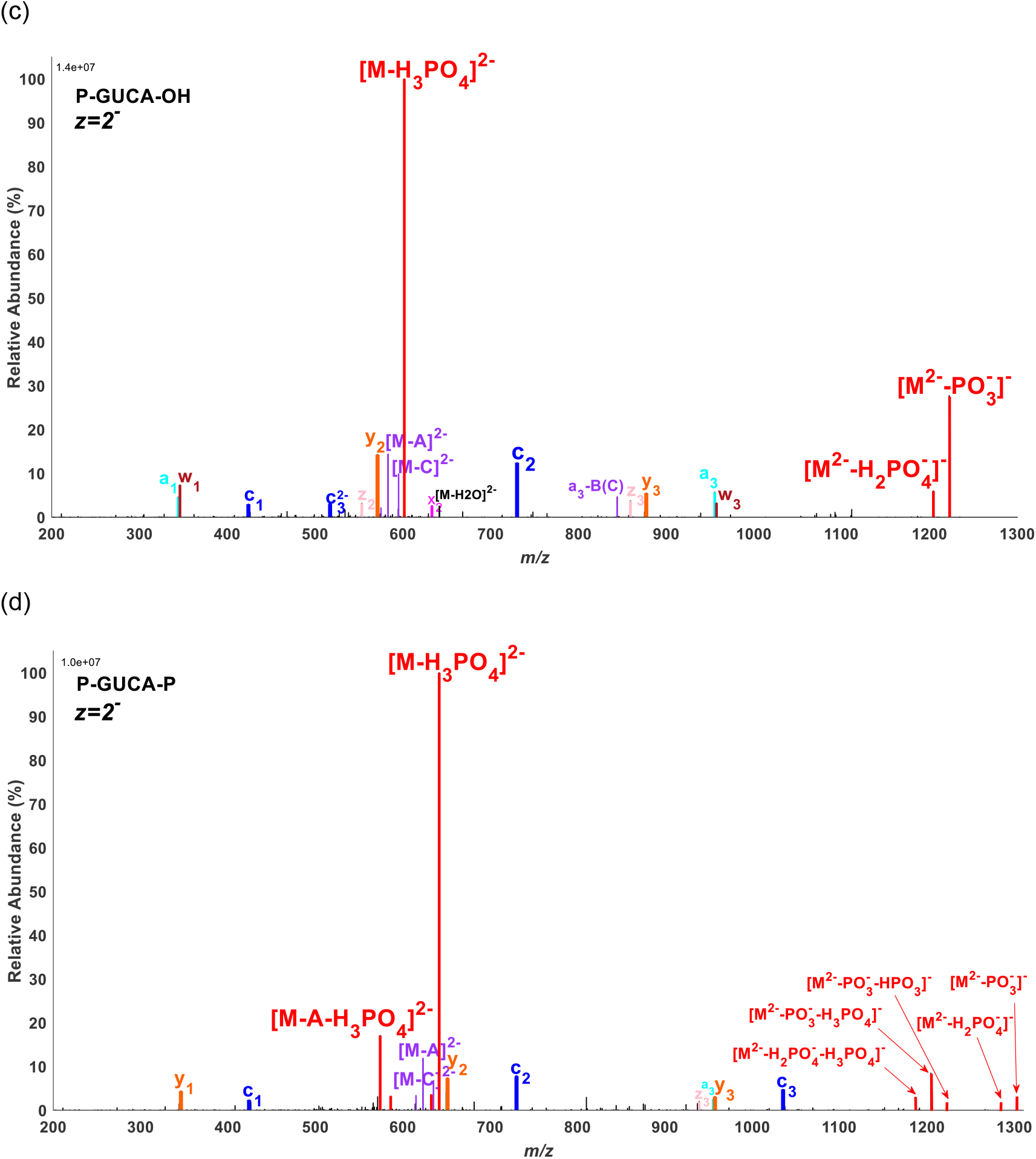
The effect of the terminal phosphate group on the CID spectra of GUCA (charge 2^-^, NCE 35%). (a) HO-GUCA-OH, (b) HO-GUCA-P, (c) P-GUCA-OH, (d) P-GUCA-P. The peaks assigned to precursor ions with a phosphoric acid loss (-H_3_PO_4_ or -H_2_PO_4_^-^) or meta-phosphoric acid loss (-HPO_3_ or -PO_3_^-^) or their combinations are annotated in red. All green peaks with an ‘i’ letter in Figure 2a are the internal fragments, generated from the cleavages of two different phosphodiester bonds.

**Figure 3.**
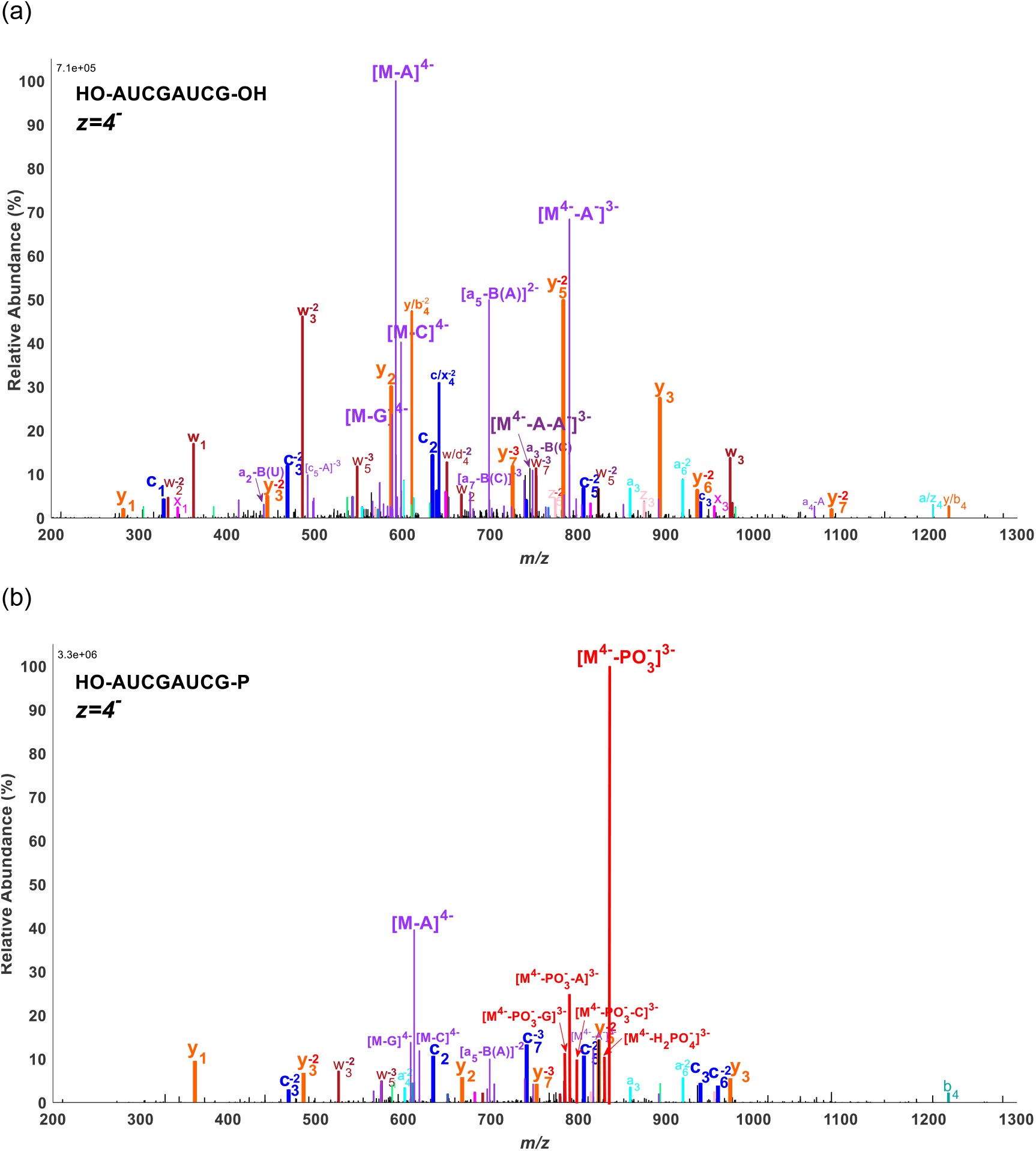
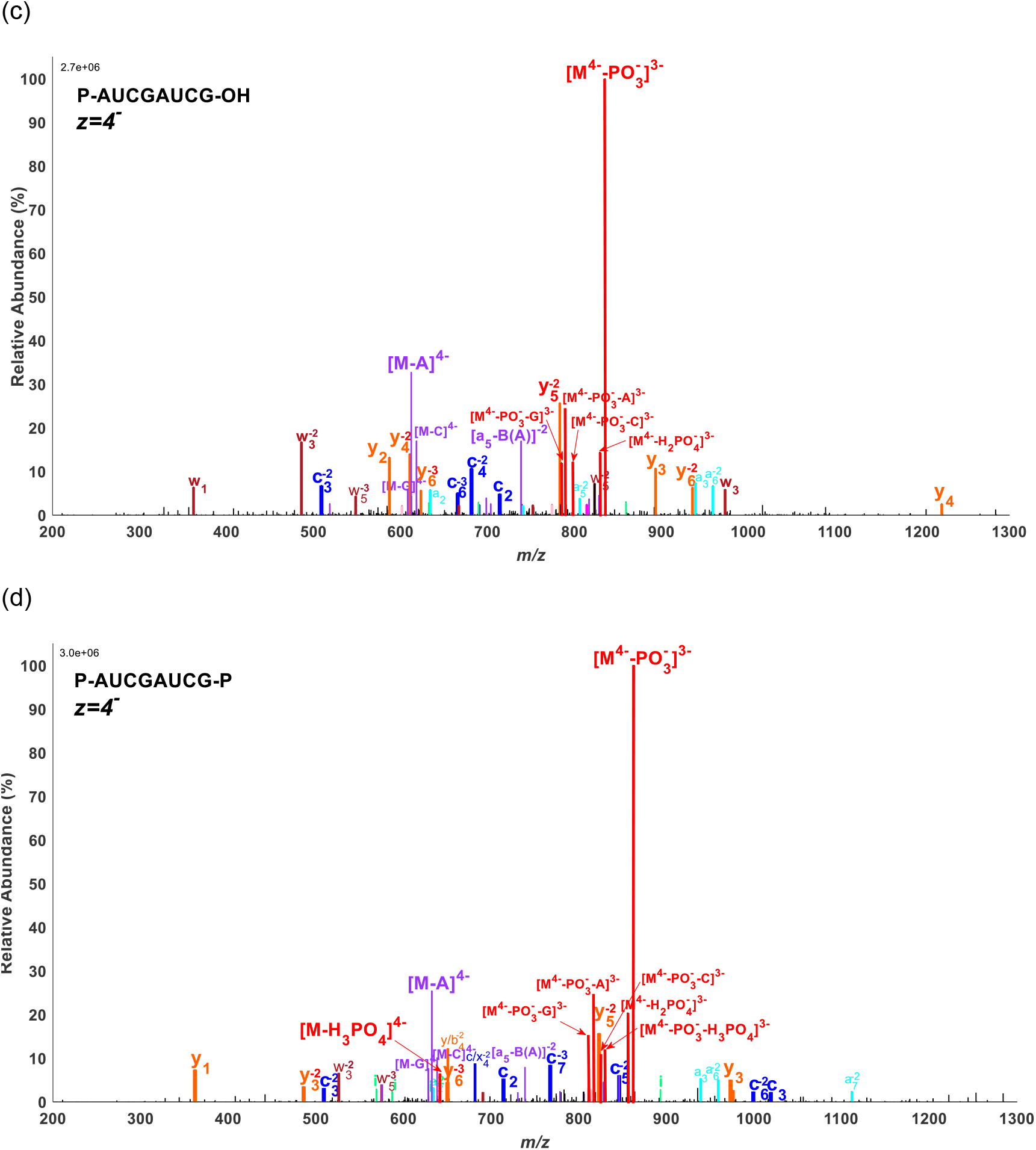
The effect of the terminal phosphate group on the CID spectra of AUCGAUCG (charge 4^-^, NCE 35%). (a) HO-AUCGAUCG-OH, (b) HO-AUCGAUCG-P, (c) P-AUCGAUCG-OH, (d) P-AUCGAUCG-P. The peaks assigned to precursor ions with a phosphoric acid loss (-H_3_PO_4_ or -H_2_PO_4_^-^) or meta-phosphoric acid loss (-HPO_3_ or -PO_3_^-^) or their combinations are annotated in red.

Additionally, we analyzed the effect of terminal phosphate groups with respect to the precursor charge state. Figure 4 shows the fractional intensity of all the sequencing ions (SI) and the fractional intensity of all the PL ions, [M-H_3_PO_4_/H_2_PO_4_^-^/HPO_3_/PO_3_^-^ or their combinations]. The fractional intensity was calculated by dividing the summed intensity of the indicated ion type with the summed intensities of all the ions in the same spectrum. For the three GUCA oligos with one or two terminal phosphate groups, precursors of charge 3^-^ generated PL ions to the highest level (52.5∼95.0%) and those of 2^-^ or 4^-^ often followed closely behind (47.1∼63.0% or 16.0∼76.5%). Likewise, for the three AUCGAUCG oligos with one or two terminal phosphate groups, precursors of charge 4^-^ and 5^-^ generated more PL ions (29.3∼55.3%) than those of 3^-^ or 6^-^ (10.9∼20.9%) and precursors of the lowest (2^-^) or the highest (7^-^) charge state appeared to have generated the least amount of PL ions (0.9∼11.6%). Briefly, loss of a phosphoric acid neutral or a metaphosphoric acid anion is the most conspicuous for precursor ions of an intermediate charge state.

**Figure 4.**
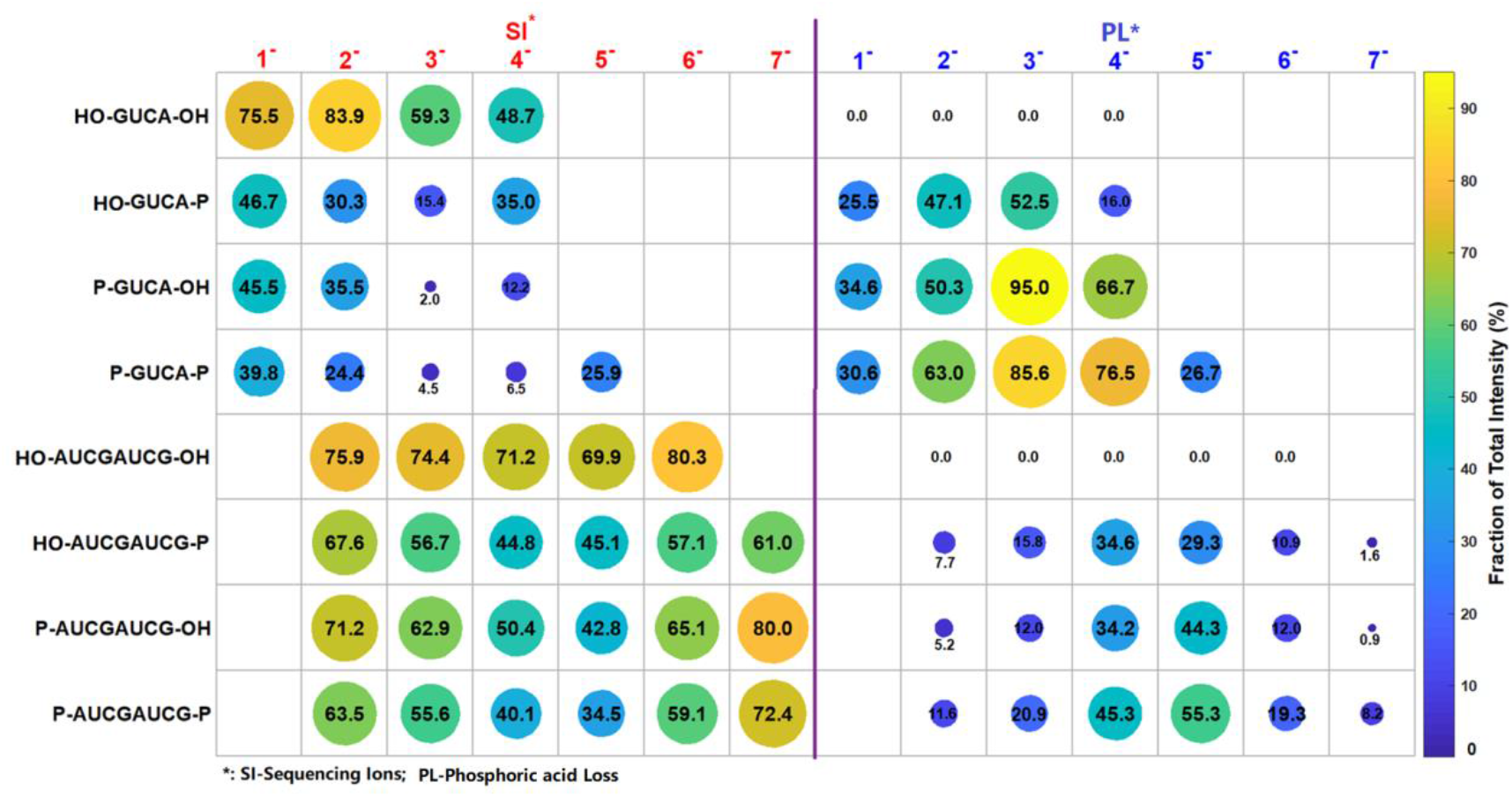
The fractional intensity of all the sequencing ions (SI) and the fractional intensity of all the (meta)phosphoric acid loss (PL) ions ([M-H_3_PO_4_/H_2_PO_4_^-^/HPO_3_/PO_3_^-^ or their combinations]). The fractional intensity (%), shown inside or below the circle, was calculated by dividing the summed intensity of the indicated ion type with the summed intensities of all the ions in the same spectrum.

### 3.3. The dehydrated pyrophosphate anion (*m/z* 158.925) in HCD spectra of RNA oligos is markedly increased by the presence of a terminal phosphate group

Lastly, we analyzed the low *m/z* ions in the HCD spectra of RNA oligos. Figure 5 shows part of the HCD spectra of the GUCA oligos (charge 2^-^) under NCE 40% (*m/z* 100∼700). The peak of *m/z* 158.93 (highlighted in red in Figure 5), was not reported in the literatures concerning MS fragmentation of RNA or DNA. A search of the NIST (National Institute of Standards and Technology) MS database of small molecules finds that this peak is frequently observed in the negative-ion HCD spectra of many molecules with multiple terminal phosphate groups, including Uridine 5’-triphosphate, phytic acid, and others. By the accurate mass analysis, we identified this peak, of *m/z* 158.925, as a dehydrated pyrophosphate anion, [HP_2_O ]^-^ Abundant pyrophosphate anions are found in MS studies of lipid A isolated from bacteria^24^ and other small molecules,^25^ which have the same terminal phosphate groups with RNA oligos. This hints that the generation of the peak of *m/z* 158.925 is more related to the terminal phosphate than the internal phosphate of RNA oligos. We will discuss it in later section.

**Figure 5.**
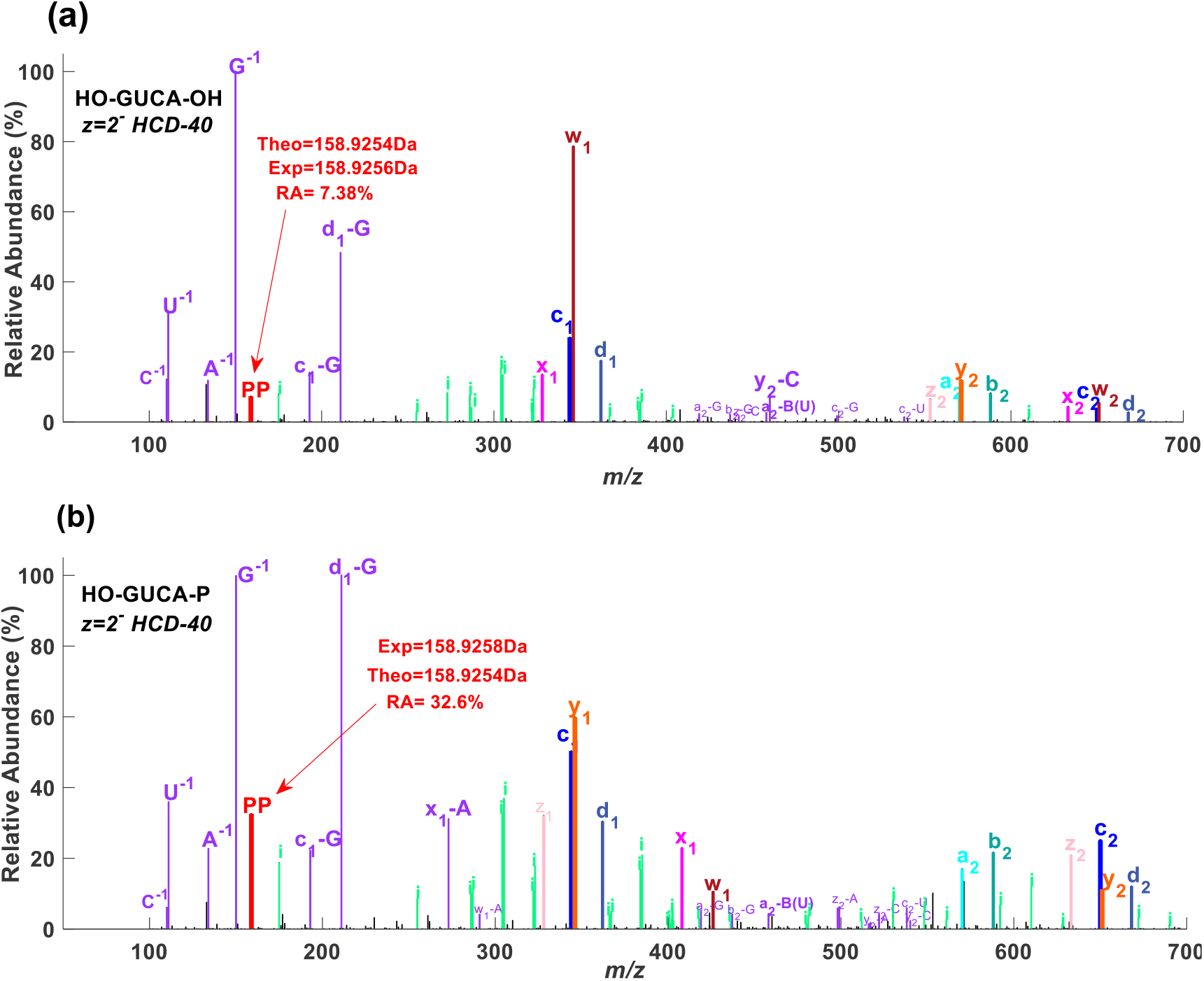
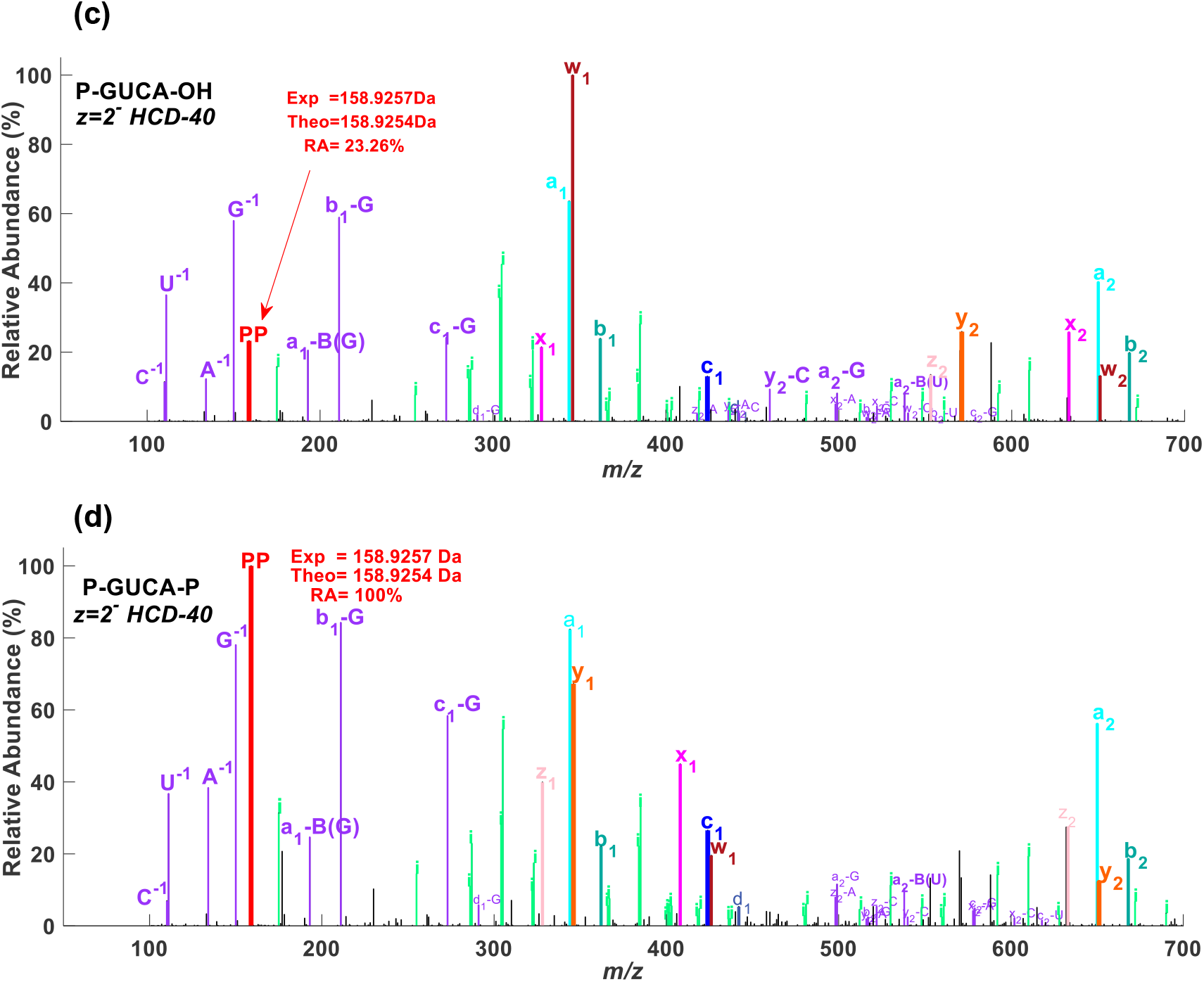
The dehydrated pyrophosphate anion (PP) in the HCD spectra of GUCA oligos (NCE 40%, charge 2^-^). (a) HO-GUCA-OH, (b) HO-GUCA-P, (c) P-GUCA-OH, (d) P-GUCA-P. All green peaks with an ‘i’ letter are the internal fragments, generated from cleavages of two different phosphodiester bonds.

In Figure 5, for simplicity, we use ‘PP’ to denote this dehydrated pyrophosphate anion, *m/z* 158.925. The relative abundance (RA) of PP increases from 7.38% for HO-GUCA-OH to 32.60% or 23.26% for HO-GUCA-P or P-GUCA-OH, respectively, and lastly to 100% for P-GUCA-P. Therefore, as the number of terminal phosphate groups increases, the RA of PP increases.

Next, we tried to find out to what extent the terminal phosphate group contributes to the formation of PP. Figure 6 shows the RA of PP as a function of NCE for eight RNA oligos. Among the four 4-nt RNA oligos (GUCA), the one with two terminal phosphate groups (P-GUCA-P) produced a much higher peak of PP than the two with only one terminal phosphate group (HO-GUCA-P and P-GUCA-OH) across NCE 25∼50. The RA of PP in the HCD spectra of P-GUCA-P reached 100% at NCE 40 or more, that is, PP became the base peak or the highest peak in those spectra. The HCD spectra of the two 4-nt RNA oligos (GUCA) with only one terminal phosphate group in turn have a higher peak of PP than those without a terminal phosphate group. The two 6-nt oligos without a terminal phosphate group,^21^ although have the same number of phosphate groups as P-GUCA-P, have in their HCD spectra a much lower peak of PP. These data show that the addition of a terminal phosphate group to an RNA oligo, boosts the formation of PP, more than the addition of an internal phosphate group, in the negative-ion HCD.

**Figure 6.**
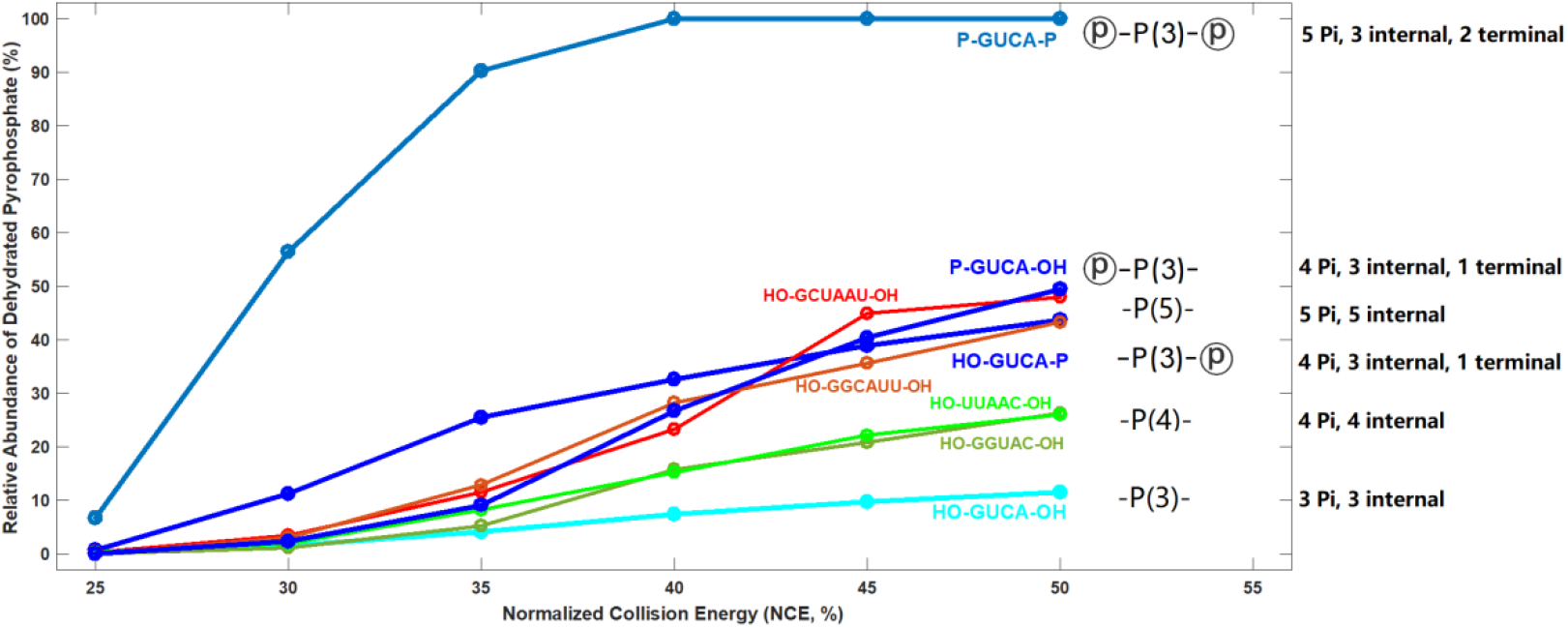
The relative abundance change of *m/z* 158.925 (dehydrated pyrophosphate anion) as a function of increasing NCE of HCD. The number of total, terminal, and internal phosphate groups are indicated.

### 3.4. CIP treatment can effectively remove terminal phosphate groups of RNA oligos

Since the presence of terminal phosphate groups decreases the intensity of sequencing ions (*a-, a-B, b-, c-, d-, w-, x-, y-, z-*ions) of RNA oligos, they should be removed for sequencing by CID or HCD. Krivos et al. used bacterial alkaline phosphatase (BAP) to remove the 3’-terminal phosphate from RNase digestion products to improve RNA sequence coverage.^26^ Here we used calf intestinal alkaline phosphatase (CIP) because it has been reported that CIP can be digested by proteinase K or inactivated by heating more readily than BAP.^27^ In other words, CIP can be removed more easily before MS analysis of the treated RNA samples, if needed.

To assess the effectiveness of CIP treatment in removing the terminal phosphate groups, especially the 5’-phosphate group, we compared the MS1 spectra of six RNA oligos with 5’- or 3’-phosphate, or both, before and after CIP treatment. As shown in Figure 7, although a fraction of the HO-X-P, P-X-OH, and P-X-P oligos (X=GUCA or AUCGAUCG) had already lost one or two phosphate groups before CIP treatment, CIP treatment clearly removed both 5’- and 3’-phosphate, and only HO-X-OH oligos were present after CIP treatment. Therefore, we recommend CIP treatment prior to MS analysis when the digested RNA samples contain either terminal phosphate group.

**Figure 7.**
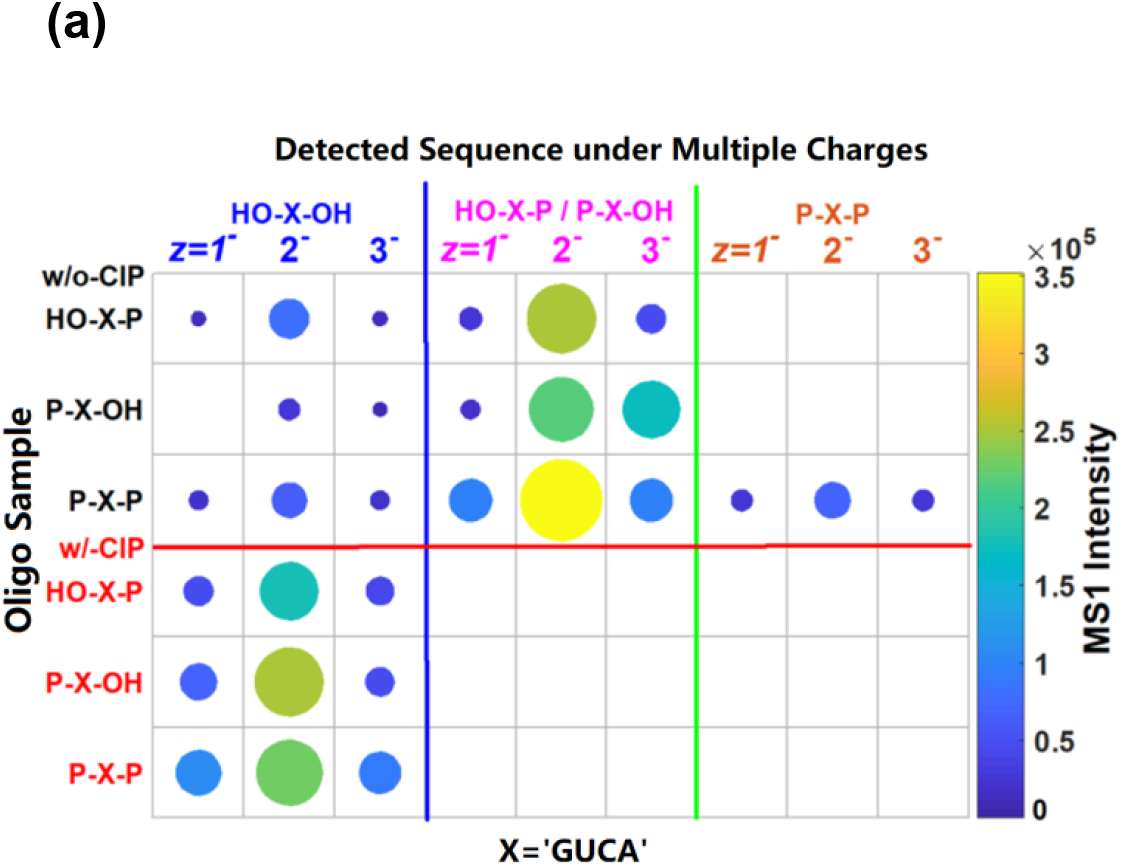
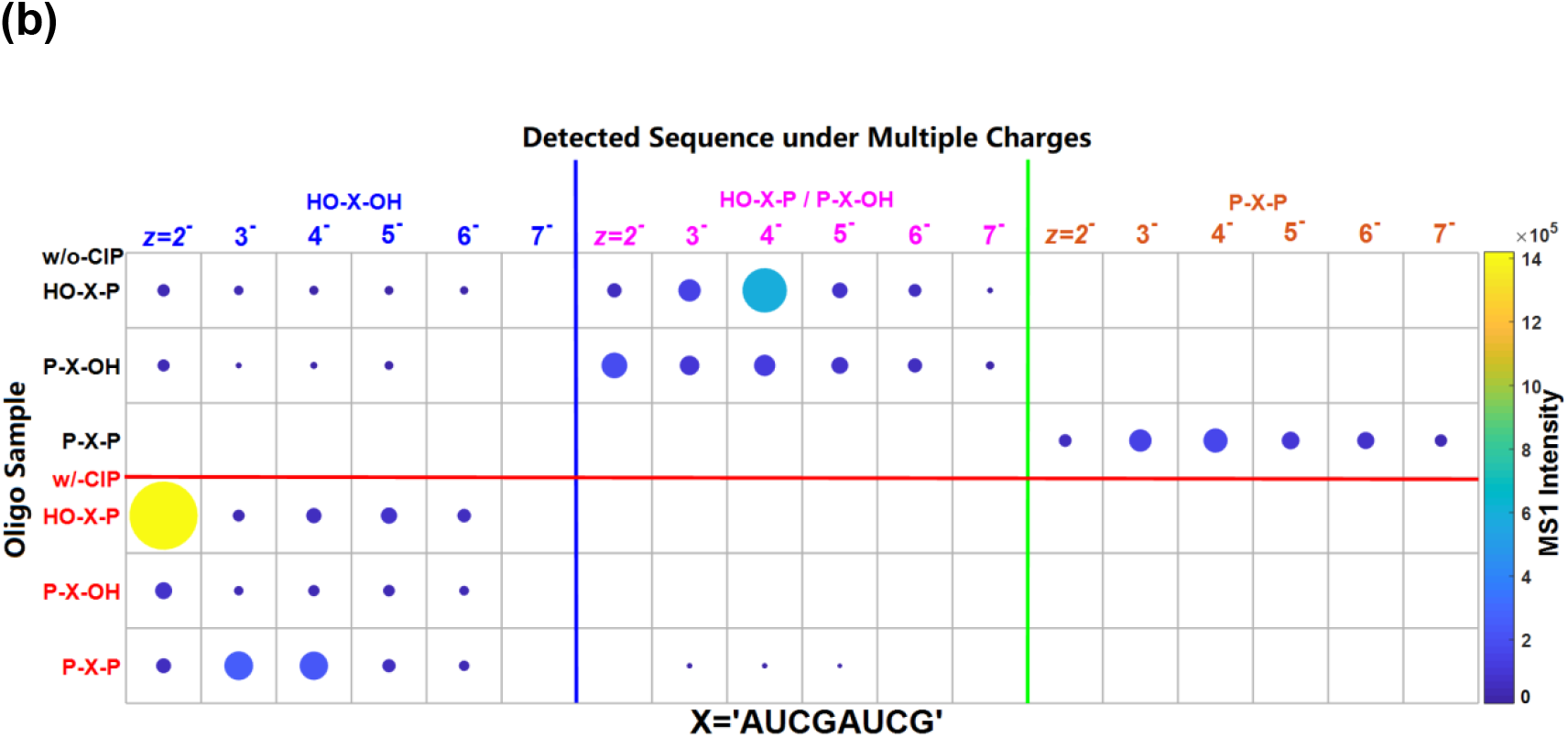
Effect of CIP treatment on the charge state and MS1 intensity of the precursors ions of RNA oligos. (a) X=‘GUCA’, (b) X=‘AUCGAUCG’. The three rows above or below the red line correspond to, respectively, the oligos before or after CIP treatment.

## 4. Conclusion and Discussions

Mass spectrometry on RNA oligonucleotides (oligos) is fundamental for understanding their structure and function. This study interrogates the influence of RNA terminal phosphate groups on the negative-ion mode collision-induced dissociation (CID) and higher-energy collisional dissociation (HCD) fragmentation. Specifically, RNA oligos from two sequences (GUCA and AUCGAUCG) and varied terminal groups (5’-OH and 3’-OH, 5’-OH and 3’-P, 5’-P and 3’-OH, 5’-P and 3’-P) were characterized by CID or HCD. Our study finds that the terminal phosphate groups have a well pronounced effect on RNA oligonucleotide fragmentation by CID or HCD, and this effect depends on the precursor charge state and the oligo length. Higher negative charges are found on precursor ions of RNA oligos with terminal phosphate groups. Loss of phosphoric acid or metaphosphoric acid neutrals or anions from precursor ions is prevalent, especially for RNA oligos with 5’- or 3’-phosphate and of medium charge states. This type of loss, neutral or charged, decreased the intensity of sequencing ions in RNA oligos with a terminal phosphate group(s). Removal of the terminal phosphate groups during sample preparation using CIP was demonstrated to be effective. In summary, this investigation on the effect of the terminal phosphate groups on CID or HCD spectra contributes to optimizing RNA mass spectrometry protocols and the development of associated software tools for high-throughput RNA analysis by CID or HCD.

As for the effect of terminal phosphate groups on the CID spectra of RNA oligos, Figure 4 shows that the predominant PL ions were observed with medium charges for both GUCA and AUCGAUCG oligos. The decrease of the fractional intensity of PL ions at higher charges, 4^-^ or 5^-^ for GUCA, and 6^-^ or 7^-^ for AUCGAUCG, can be partially attributed to the increase of nucleobase losses from precursor ions, which are dominant and competitive with the formation of PL ions at higher charge states. The nucleobase losses from precursor ions for RNA oligos with one or two phosphate groups have the similar patterns with RNA oligos with both 5’- and 3’-OH termini. The latter has been studied in detail in ref. 21.

How the dehydrated pyrophosphate anion (*m/z* 158.925) is generated awaits future investigation. Thomas et al.^25^ studied MS fragmentation of phospholipid headgroups and proposed a scheme that involves rearrangement of two terminal phosphate groups from two different ions into a pyrophosphate anion. It would be interesting to see whether a similar mechanism can explain the formation of dehydrated pyrophosphate from RNA oligos in the gas phase.

## Acknowledgments

This study was supported by the National Key Research and Development Program of China (Grant 2020YFF01014505), the Ministry of Science and Technology of China, and Beijing Municipal Science and Technology Commission. We also thank Yan Ma in Metabolomics Center, National Institute of Biological Sciences, Beijing, China (NIBS) and Ji-Shuai Zhang in the Department of Chemistry, Peking University, China for their constructive advice concerning the identification of the peak at *m/z* 158.925 in the HCD spectra.

## Associated Content

### 1. Supplemental Table S1

The detailed information of eight RNA oligonucleotides synthesized in this manuscript

### 2. Supplemental Figure S1

The MS1 signal intensity of eight RNA oligonucleotides.

