## Supplemental Figure S1 for "Effect of terminal phosphate groups on collisional dissociation of RNA oligonucleotide anions"

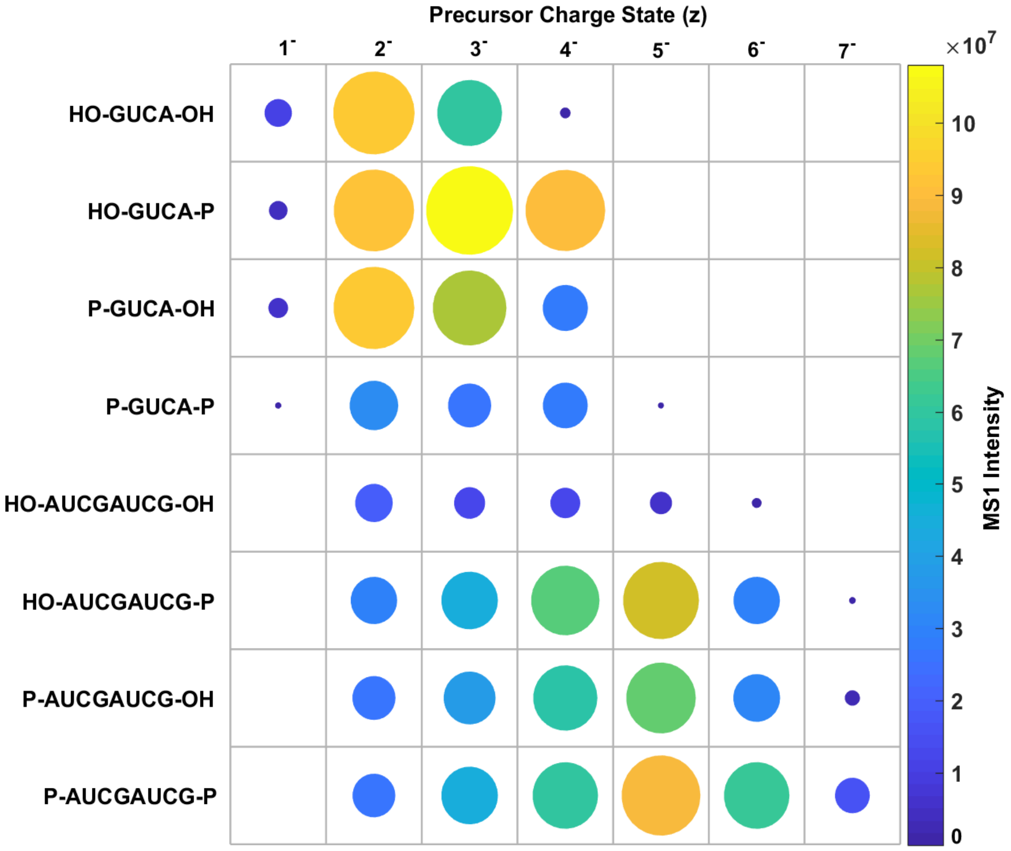


MS1 signal intensity of eight RNA oligonucleotides. A higher intensity is denoted by a larger diameter and a warmer color.
